# Anterior insula activity increased by cued risky wins in healthy volunteers

**DOI:** 10.64898/2026.06.30.735658

**Authors:** Lester C Tong, Brandon J. Forys, Claire A. Hales, Luke Clark, Catharine A. Winstanley

**Affiliations:** Department of Psychology, Djavad Mowafaghian Centre for Brain Health, University of British Columbia, Vancouver, BC, Canada

## Abstract

Audiovisual cues (‘bells and whistles’) are ubiquitous in commercial gambling products. Pairing wins with sound and light cues in laboratory-based gambling paradigms increases risky choice, but the neurocognitive basis of this effect is unclear. Here we compared patterns of neural activation using functional MRI in healthy volunteers (n = 31) while they performed a two-choice lottery task. Reward-paired cues were either present or absent in a mixed-block, event-related design. As predicted, participants made riskier choices on cued trials. Choice latencies were also longer when cues were present, particularly on trials following a win. Activity within the nucleus accumbens and orbitofrontal cortex was greater during the decision phase when participants made risky choices. Nucleus accumbens signal was also greater when participants were anticipating risky outcomes, and in response to risky wins. Contrary to our pre-registered hypotheses, cue condition did not alter patterns of activity across any task phase, in either of these *a priori* regions of interest. As such, cue-induced risky choice does not appear to be driven by altered representation of risk or value within this canonically reward-sensitive circuitry. Instead, exploratory analyses revealed that the anterior insula was selectively activated by cued, risky wins. Such activation may signal the saliency of these events, or their emotional impact, and may reflect one mechanism through which cue-induced craving develops in vulnerable individuals.

## Introduction

Gambling is a real-world example of risky decision-making with a recognized capacity for harm, and this has led many countries to adopt a public health perspective on gambling that includes consideration of addictive design features (Wardle et al., 2024). Sound and light stimulation (adding ‘bells and whistles’) is ubiquitous in modern slot machines, other electronic gambling machines (EGMs), and online gambling products, to signal and augment winning outcomes (Barrus et al., 2016; Newall, 2025). Video game developers seeking to increase product engagement are also increasingly embedding the structural characteristics of slot machines within their designs (Flayelle et al., 2023). Pairing audiovisual cues with reward delivery increases risky choice in healthy volunteers (Cherkasova et al., 2018; Spetch et al., 2020), and in rats performing the rat gambling task (rGT)(Adams et al., 2017; Barrus & Winstanley, 2016; Langdon et al., 2019). Determining the neural mechanisms through which win-paired audiovisual cues promote risky choice may help us understand why EGMs and other heavily-cued games can be so addictive.

Elevated risky decision-making has been reliably associated with both gambling and substance addictions, and these conditions further occur comorbidly (Brand et al., 2025; Dowling et al., 2015; Kovacs et al., 2017; Petry, 2000, 2001). In rodents, repeated engagement with highly cued and/or probabilistic reward schedules amplifies the reinforcing effects of cocaine and amphetamine, suggesting that pairing wins with sensory stimulation may drive a pro-addiction phenotype (Ferland et al., 2019; Hynes et al., 2023; Mascia et al., 2018; Singer et al., 2012; Zack et al., 2014). Heightened cue reactivity also drives craving across addictive disorders (Carter & Tiffany, 1999; Noori et al., 2016; Starcke et al., 2018), and in people with disordered gambling, attentional bias towards gambling-related cues facilitates risky decision making and correlates with gambling severity (Ciccarelli et al., 2020).

Cue-driven risky choice does not appear to reflect a predisposition to sign- or goal-tracking in humans (Cherkasova et al., 2024), while rodent data suggest a relationship only when an operant lever is used as the sign, rather than an audiovisual stimulus (Ferland et al., 2019; Swintosky et al., 2021). Win-concurrent cues therefore appear to work through a mechanism distinct from pure Pavlovian approach (see (Heck et al., 2026) for recent review of this literature). During laboratory-based gambling tasks, people spend less time looking at information signaling the probability of winning when reward-paired cues are included in-game (Baumann et al., 2025; Cherkasova et al., 2018), suggesting cues may cause participants to develop an inaccurate model of the decision space. Rodent data suggest that decision-making patterns are less sensitive to reinforcer devaluation when cues promote risky choice (Hathaway et al., 2022; Hathaway et al., 2021), indicating decision making is not truly goal-directed (i.e. model-based) under such conditions. The orbitofrontal cortex (OFC) is essential for the goal-directed behavior of cue-guided choice in non-human primates (Baxter et al., 2000), and neuronal activity in this region represents subjective outcome value in multiple species (Gardner et al., 2020; Gottfried et al., 2003; Howard & Kahnt, 2017; Kuwabara et al., 2020; Padoa-Schioppa & Assad, 2006). Infusing a serotonin 2C antagonist directly into the lateral OFC of rats decreased risky choice on the cued rGT and restored sensitivity to reinforcer devaluation, such that that win-paired cues may alter OFC functioning to both reduce goal-directed control and enhance risky decision-making (Hathaway et al., 2021).

The ventromedial prefrontal cortex (vmPFC, which incorporates the OFC) and the ventral striatum are strongly activated by rewarding outcomes in fMRI studies (Bartra et al., 2013; Lopez-Gamundi et al., 2021). Both *ex vivo* cFos imaging in rats that had performed the cued rGT (Mortazavi et al., 2023), and fMRI connectivity analyses (Limbrick-Oldfield et al., 2017; Park et al., 2010) indicate strongly correlated activity between these regions. Reward-predictive cues enhance dopamine release within the nucleus accumbens (NAcc) (Gan et al., 2010; Lefner et al., 2022), and dampening this signal reduced risky choice in male rats performing the cued rGT (Hynes et al., 2020). We therefore pre-registered hypotheses that activity within these regions of interest (ROIs) would be more pronounced when participants perform cued vs uncued trials of a two-choice lottery task.

The conjunction of cues with reward may affect different phases of the decision process, from the evaluation of options at the time of decision, to the prediction or anticipation of outcomes, to the moment of feedback and outcome delivery (Ernst et al., 2004). fMRI research indicates that ventral striatum is particularly active during reward anticipation, and tracking reward prediction error (Hare et al., 2008; Rohe et al., 2012). vmPFC signals are known to also track reward receipt (Knutson & Cooper, 2005; Knutson & Wimmer, 2007; Yacubian et al., 2006), and co-activation of both regions is implicated in the computation of value during decision evaluation (Clithero & Rangel, 2014). Despite this potential overlap in neural circuitry recruited across task phases, activity in the OFC and dorsal striatum was specifically enhanced in individuals with problem gambling during reward anticipation (van Holst et al., 2012). Conversely, the ventral striatum and vmPFC were relatively underactive when probabilistic rewards were delivered in individuals classified as having pathological gambling (gambling disorder), and activity in both regions was negatively correlated with a measure of gambling severity (Reuter et al., 2005). We therefore evaluated activity within our ROIs across time of decision, anticipation, and feedback phases of the task, but remained agnostic as to whether activity would be elevated or dampened in the cued relative to uncued condition.

Reinforcement learning algorithms model the assumption that mental representations of option values are updated on a trial-by-trial basis in response to positive and negative outcomes, and used to guide decision making. Somewhat contrary to the hypothesis that combining rewards with audiovisual cues will potentiate activity in the reward system, RL models did not find evidence of greater learning from rewards on the cued rGT (Hales, Hwang, et al., 2025; Langdon et al., 2019). Instead, option value was not updated as much when penalties occurred. Such analyses suggest that cue-induced risky choice arises from hyposensitivity to negative outcomes, rather than hypersensitivity to rewards. A meta-analysis of neuroimaging studies using RL modelling found that learning from penalties is associated with activity in the anterior insula (AntIns) (Garrison et al., 2013). RL models have also linked neural responses in AntIns to poorer learning from punishments in individuals with gambling disorder (Iigaya et al., 2025; Suzuki et al., 2023) as well as conduct disorder (Elster et al., 2025). The AntIns was therefore included as a further exploratory ROI in our analyses.

## Methods

### Participants

A sample of 31 healthy volunteers was recruited primarily from University of British Columbia mailing lists, including the psychology department ‘human subjects pool’ and paid subjects list. Participants were 11 male and 20 female, aged 19-44. Prior to scanning, participants were phone screened for eligibility, including MRI contra-indications, and administration of the problem gambling severity index (PGSI). Participants scoring PGSI 2+ or who reported any history of gambling disorder, past enrolment in gambling self-exclusion programs, or self-reported history of substance use, were excluded. Participants were required to have normal or corrected-to-normal vision and hearing. The study was conducted in accordance with institutional guidelines and the Declaration of Helsinki, and was approved by the Research Ethics Board of the University of British Columbia. Participants gave written informed consent.

### Vancouver Gambling Task

To test how audiovisual cues modulate risky choice and reward-related brain activity, we modified a two-choice lottery task, labelled the Vancouver Gambling Task (VGT), to enable a mixed-block, event-related fMRI design. Previous studies using the VGT have presented ‘cued’ and ‘uncued’ gambles in separate blocks of the task (Cherkasova et al., 2018; Cherkasova et al., 2024). By using a mixed-block design, we can assess transient trial-by-trial effects, modelling both trial difficulty (as the difference in expected value between the pair of gambles presented) and the outcome of the prior trial (Limbrick-Oldfield et al., 2020), as well as more sustained effects of time. This design also minimizes the potential for order effects, while making the task more dynamic and engaging.

The VGT was completed in a single MRI session, comprising two fMRI sequences with a short break in between them. Participants were presented with 68 pairs of gain-only gambles in each sequence, for a total of 136 trials. In each trial, participants chose between an option with a high chance of winning a small amount (e.g., 70% chance of winning $2, henceforth the low risk option) versus a smaller chance of winning a larger amount (e.g., 20% chance of winning $5, henceforth the high risk option) (Fig 1). The winning probabilities are represented by the shaded area in each pie chart, while the reward size was presented underneath each pie chart. In order to elicit a range of both risk-seeking and risk-averse responses from each participant, the expected values of the two options vary such that on some trials, the high-risk option is the mathematically rational choice, whereas on other trials the low-risk option is optimal. The task is incentive compatible, with points accumulated translating directly into a monetary bonus on completion ($0.10 CAD per point).

**Figure 1.**
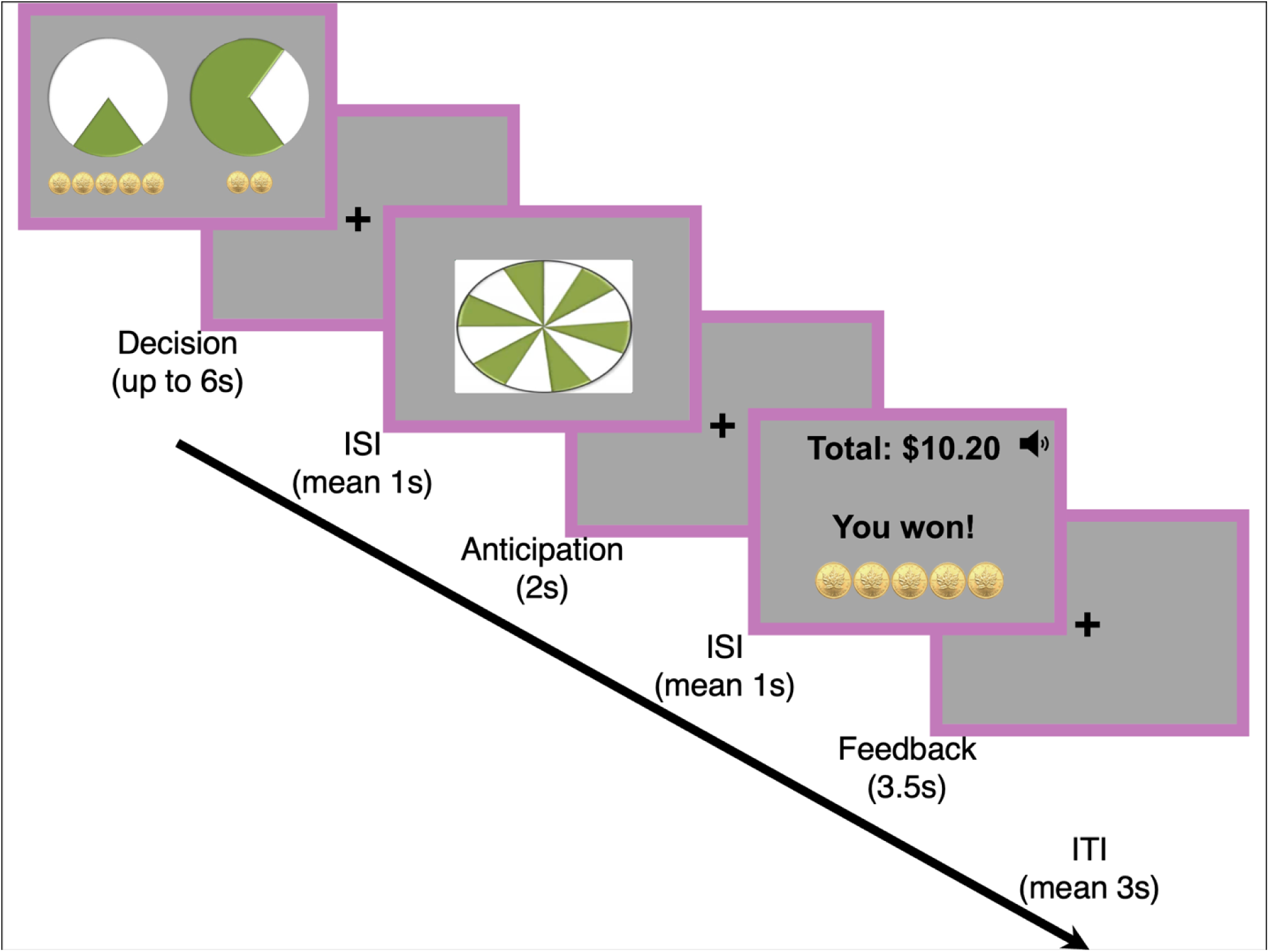
Vancouver Gambling Task. In the decision phase, participants chose between two gambles, with probability represented as the green area of a pie chart, and magnitude indicated underneath either as a number of coins (cued) or an Arabic numeral (uncued). In the anticipation phase, participants viewed a green and white spinning wheel while uncertain of the trial outcome. Participants viewed the outcome of their decision in the feedback phase, either as animated coins and auditory cues (cued), or just as an Arabic numeral (uncued). Non wins simply displayed the participant’s running total accompanied by the text “No win…”

On each trial, the participant either won the reward depicted in the chosen gamble or received nothing: no losses were presented in this version of the task. On uncued trials, rewarding outcomes were displayed only in plain text (e.g. “You won! $5”). On the cued trials, in addition to the text “You won!”, rewarding outcomes were displayed as a number of coins, accompanied by further audiovisual cues that scaled in intensity and complexity with the magnitude of the win (Cherkasova et al., 2018). The visual enhancement was as follows: 1 token was represented as a static two-dimensional image of 1 gold coin; 2 tokens as 2 static two-dimensional gold coins with a sparkle (static luminance enhancement); 3 tokens as 3 static three-dimensional gold coins with 3 sparkles (static luminance and depth enhancement); 4 tokens as 4 three-dimensional gold coins with a sparkle running along the circumference of each coin (dynamic luminance and depth enhancement); 5 tokens as 5 three-dimensional spinning gold coins with a sparkle running along the circumference of each coin (dynamic luminance, depth and motion enhancement). Despite the visual enhancement, the visual stimuli for the different reward magnitudes were not substantially different in terms of the overall average luminance (1 coin: 100.79 cd/m^2^; 2 coins: 101.54 cd/m^2^; 3 coins: 98.93 cd/m^2^; 4 coins: 94.37 cd/m^2^; 5 coins: 101.18 cd/m^2^) and color of the image (1 coin: .215 v’, .500 u’; 2 coins: .240 v’, .531 u’; 3 coins: .228 v’, .525 u’; 4 coins: .186 v’, .387u’; 5 coins: .229 v’, .528 u’); the 3-dimensional images were slightly lower in luminance because of the shading. The average luminance of the uncued feedback image was 102.44 cd/m^2^. The auditory enhancement consisted of sounds taken from a casino library and edited to conform to the temporal structure of the task. The tunes accompanying the rewards progressively increased in duration (1200 to 2700 ms), loudness (44 to 52dB) and complexity (variation in tempo and pitch) as the reward magnitude increased from 1 token to 5 tokens. Prior to the scan, participants were instructed regarding the task and the cued and uncued conditions, and practiced the task for five minutes.

Trials were presented in one of two pseudorandomized orders, with alternating blocks of 6 cued trials and 6 uncued trials. (In the last 2 blocks this increased to 7 trials each, for a total of 68 trials per scan). The cued vs uncued blocks were distinguished by a colored border (purple or yellow to minimize color associations, counterbalanced). A trial (Fig 1) proceeded as follows: i) fixation cross inter-trial-interval (variable duration, average 3000ms), ii) decision phase, in which participants chose between the two gambles with a button press (duration determined by response latency, about 2000ms on average, maximum of 6000ms), followed by a brief inter-stimulus-interval (variable duration, average 1000ms), iii) anticipation phase, in which the pies ‘spun’ similar to a roulette wheel (fixed 2000ms), followed by another brief inter-stimulus-interval (variable duration, average 1000ms), iv) feedback phase, in which the gamble outcome is presented, including the augmented audiovisual stimuli to winning outcomes in the cued trials, or the text ‘no win’ if they did not win (fixed duration, 3500ms). In the first two blocks, winning outcomes were forced on the first, sixth, seventh, and ninth trials to standardize exposure to the cued and uncued conditions at the start of the task. After that point, outcomes were determined randomly, in line with the displayed probability for that gamble. The timing structure with variable inter-stimulus-intervals was intended to deconvolve the decision-, anticipation-, and feedback-related fMRI responses.

All but one participant chose the risky option more frequently when the EV of the risky option was greater than the safe option. Parallel analyses excluding this participant did not change the pattern or significance of the reported results (see supplementary material). Participants responded on the vast majority of trials, timing out on only 22 trials (0.52% of 4216 total trials across all participants). No responses were faster than 300ms.

### fMRI

Participants were scanned using a 3T GE Signa PET/MR scanner with a 16-channel head coil, with whole brain coverage achieved using 50 3-mm thick axial slices (in-plane resolution = 3mm isotropic, no gap, interleaved acquisition, matrix size 86×86) extending from the mid pons to the top of the skull. During the scan set-up, the volume was calibrated to ensure participants could hear the auditory cues. The MRI scan began with the 5 minute T1-weighted anatomical scan, followed by the two, approximately 14 minute, VGT functional scans. The high-resolution T1-weighted anatomical image was acquired using a MPRAGE sequence (TR=8.4ms, TE=3.2ms, flip angle=8°, 206 slices, 1mm isotropic, matrix size 256×256). The functional scans were collected with a T2*-weighted gradient-echo pulse sequence with a multiband factor of 2 (TR=1.5s, TE=25ms, flip angle=72°).

### Behavioral Analysis

A series of generalized linear mixed models were conducted to test the effects of cue condition, the difference between the expected values of the two gambles, expressed as a ratio (henceforth EVR), and previous trial outcome, on risky choice (see Cherkasova JN 2018, Limbrick-Oldfield et al 2020). As a binary outcome variable, this used a logit link. Subjects were modeled as random intercepts, and binary regressors were effect coded.

For the corresponding analysis of decision latency, the absolute value of the EVR was used as a predictor, because regardless of sign, trials with EVR closer to zero (i.e. two gambles closer in expected value) were considered ‘more difficult’.

### Neuroimaging Analysis

fMRI preprocessing was conducted using Analysis of Functional Neural Images (AFNI) software. The first six volume acquisitions were discarded. Brain images were corrected for variation in slice-timing using sinc-interpolation, motion corrected using six-parameter affine transformations to realign each volume to the volume acquired with closest temporal proximity to the anatomical scan, and spatially smoothed with a Gaussian 4 mm fullwidth at half-maximum kernel. The resulting images were normalized to percent signal change within voxel and high-pass filtered at 1/90 Hz. Anatomical images were coregistered to the most temporally proximal functional volume, and spatially normalized by warping to the “Colin” brain template in Talairach space.

A whole-brain multivariate ANOVA was conducted in AFNI with risky choice (risk/safe), outcome (win/non-win), cue condition (cued/uncued), phase (decision, anticipation, feedback), and fMRI scan (run 1/run 2) as within subjects variables. General linear tests were specified within the same model to test for the main effects of each variable, and for interactions with cue condition.

We conducted pre-registered region-of-interest (ROI) analyses on NAcc and OFC (https://doi.org/10.17605/OSF.IO/J6SE4). We also conducted exploratory analyses on anterior insula (AntIns) which was supported by whole brain results. We used NAcc and AntIns masks publicly available with AFNI software, generated by Rutvik Desai using Destrieux, DesikanKilliany, and Freesurfer parcellations in Talairach space. For the OFC ROI, we generated a mask with Neurosynth compose (compose.neurosynth.org; https://compose.neurosynth.org/projects/RTE7PE39EyZy/meta-analyses/a6h65mLQvNH6), using search terms “gambling”, and “orbitofrontal cortex” or “ventromedial prefrontal cortex”, and we exported the resulting ventromedial prefrontal cluster, thresholded at FDR q<0.05. As manipulation checks, given that our cue condition involved both visual and auditory feedback, primary visual cortex (V1) and primary auditory cortex (A1) data were also extracted from masks using the TT_Daemon atlas distributed with AFNI software. All ROIs were warped from Talairach to subject native space by inverting the warps derived from spatial normalization. Preprocessed data were averaged within each ROI, and activity time series were extracted for computing neural activity metrics. We began by plotting within-ROI averaged activity over time for each ROI, in the decision, anticipation, and feedback phases of the task (Fig 2). Inspection of the fMRI time series in primary auditory cortex and primary visual cortex indicated that activity peaked at 3 TRs post onset. We therefore used the 3^rd^ TR to adjust for the hemodynamic lag in all subsequent analyses. Given a TR of 1.5s, this equated to approximately 4.5 seconds post stimulus onset, which corresponds to the typical 5-6 second hemodynamic lag in fMRI.

**Figure 2.**
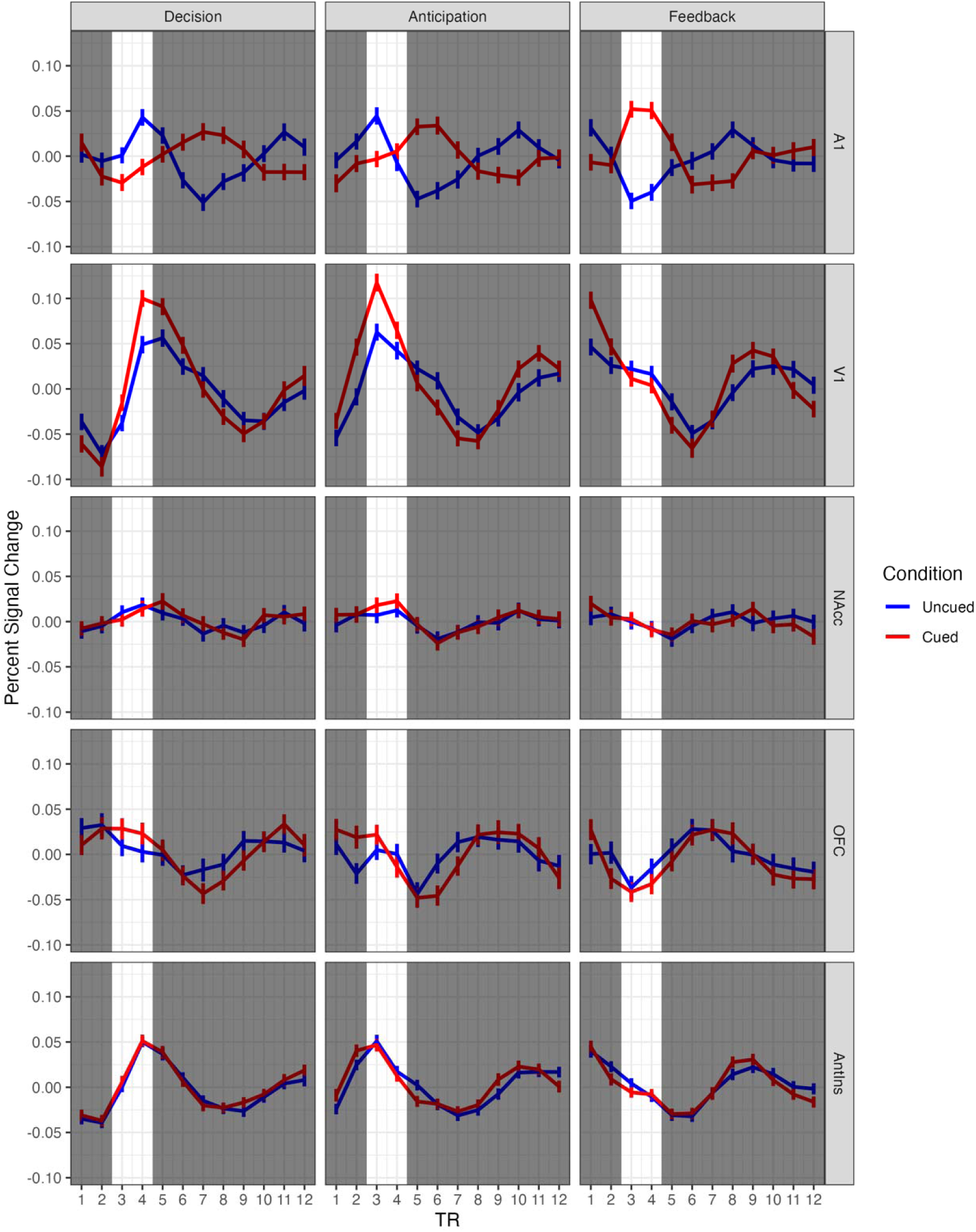
ROI time series in each VGT phase. Time series in primary auditory cortex (A1) during feedback (the only auditory stimulus), and primary visual cortex (V1) during anticipation (the most consistent visual motion) were inspected to determine the TR window (TRs 3-4, highlighted) for subsequent ROI analyses.

## Results

### Behavior on the VGT

Descriptively, participants selected the risky option on average 46.5% ± 0.5 of the time (Fig 3A), thus displaying slight overall risk aversion. All participants distributed their choices between the two options, with between 10% and 90% risky choices. As hypothesized, participants chose the risky option more often on cued compared to uncued trials (mean: 50.5% ±0.5 vs 42.4% ±0.49; t_30_ = 2.35, *p* = 0.026).

**Figure 3.**
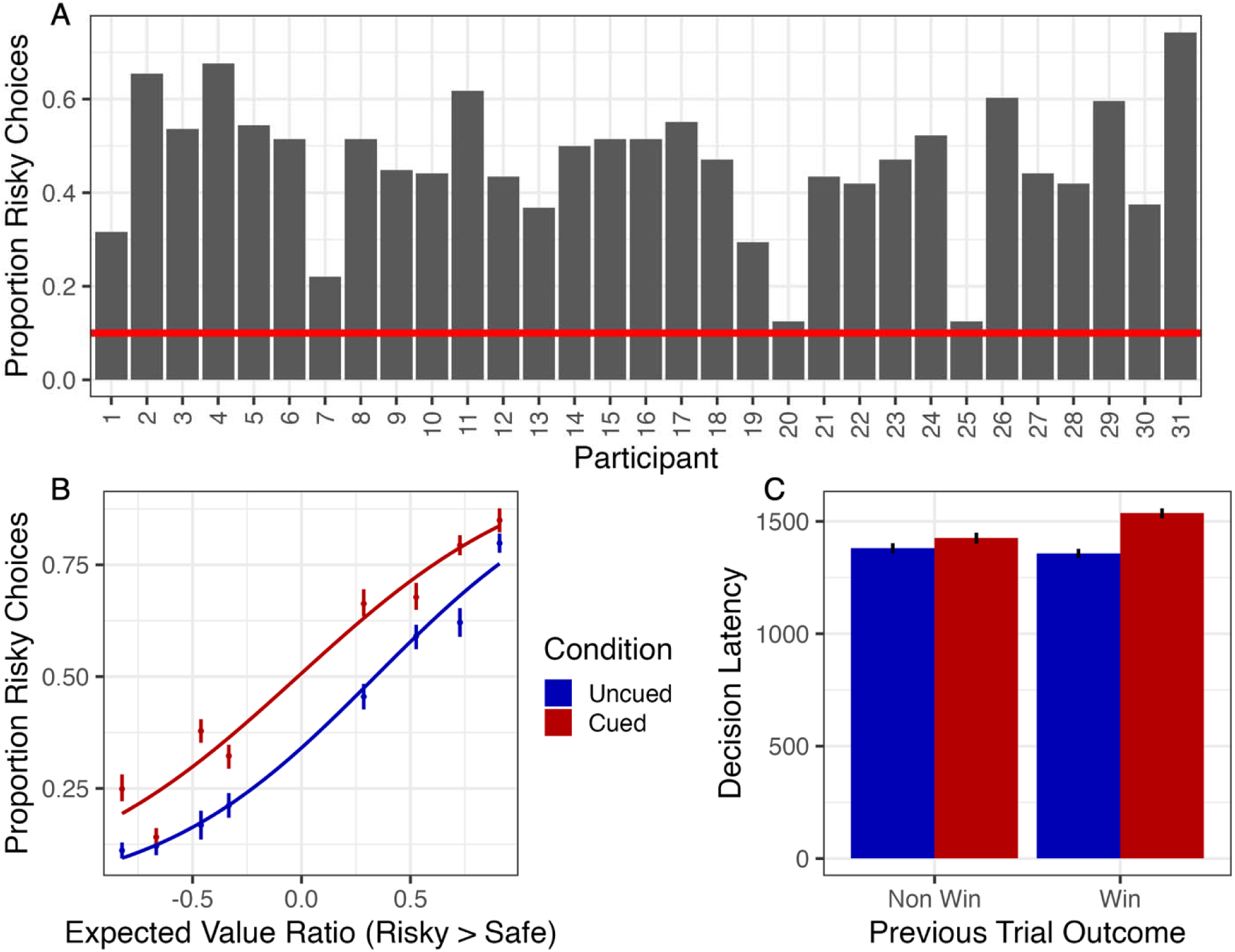
Behavioural data. (A) Proportion of risky choices in the entire VGT across individual participants. Participants varied on tendency to choose the risky gamble on average. The red line indicates the 10% threshold for excluding participants for lack of response variability. (B) Proportion of risky choices as a function of the ratio of expected value between the risky choice to the safe choice. As the EV of the risky choice increased relative to the EV of the safe choice, participants made more risky choices. Moreover, participants were more risky on average in the cued condition compared to the uncued condition at every EVR value. (C) Average decision latency was greater in cued trials than uncued trials, as well as in trials with previous wins compared to previous nonwins. A post hoc multiple regression revealed a significant interaction between cue condition and previous wins, indicating that, in the presence of cues, winning on the previous trial increased reaction time even more.

In the models for risky choice, a significant effect of the cue condition (pre-registered hypothesis 1) was confirmed, both in a simple model (b = 0.177, *p* < 0.001; Table 1), and in a multiple regression that added the EVR term and the previous trial outcome (cue condition, b = 0.374, *p* < 0.001; EVR, b = 2.10, *p* < 0.001; previous trial, b = −0.173, *p* < 0.001; Fig 3B). These coefficients indicate that participants chose risks more frequently on trials with greater EVR, and less frequently when they had just won the previous trial.

**Table 1.**
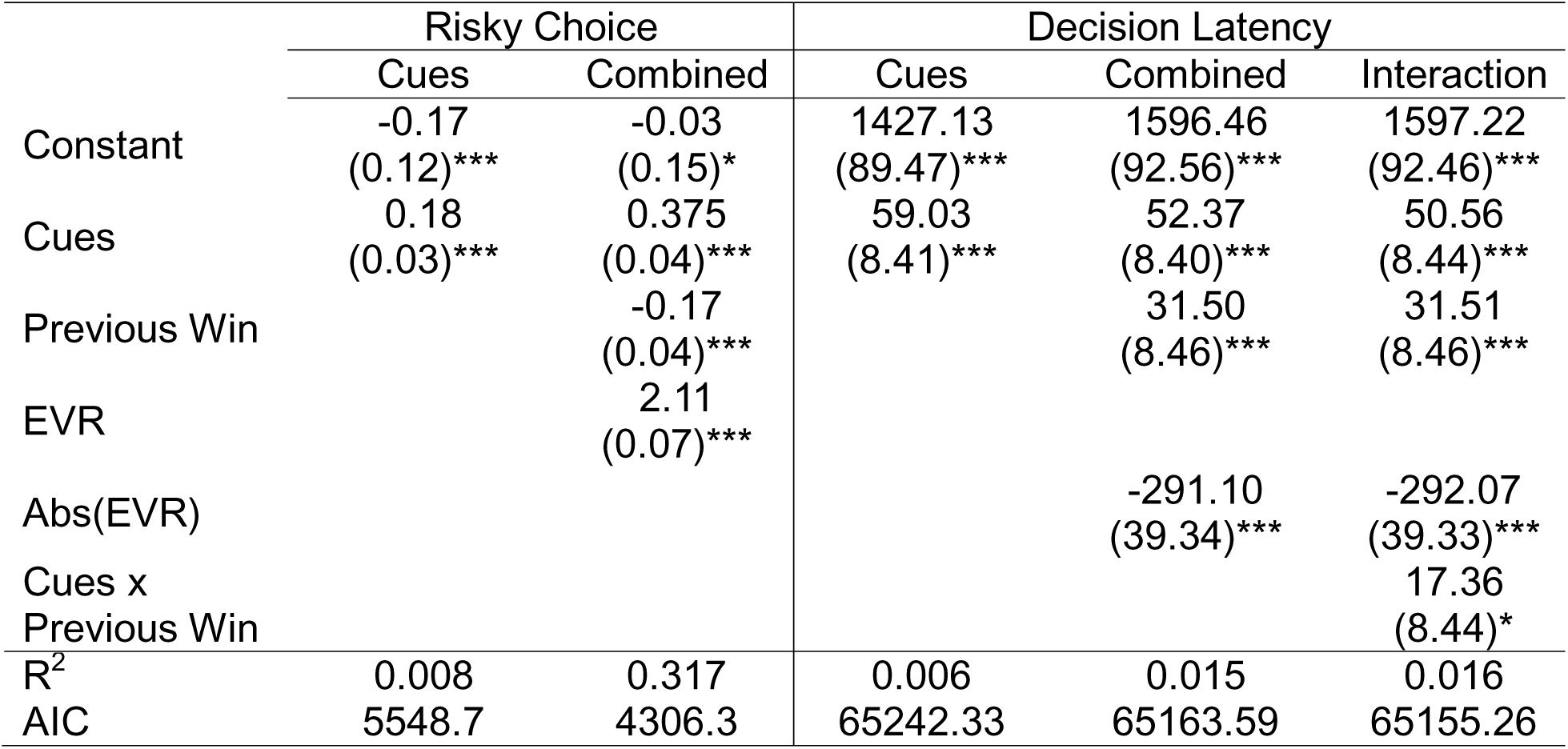
Behavioral regression results. Statistics are coefficients and SEs. Significance (two-tailed) is indicated by \**p* < 0.05; \*\**p* < 0.01; \*\*\**p* < 0.001; ^†^*p* < 0.10 (nonsignificant trend).

Participants responded in 1427.13 ± 734.52 ms on average. Uncued decision latencies (1372.47 ± 478.62) were faster on average than cued decision latencies (1496.25 ± 533.11, t_30_ = 3.92, *p* < 0.001). In the models for decision latency, cues increased decision latency by b = 59.03ms (*p* < 0.001) compared to the grand mean. Cues continued to increase decision latency when controlling for absolute EVR and previous trial outcome (b = 52.37, *p* < 0.001). As expected, absolute EVR strongly negatively predicted decision latency (b = −291.10, *p* < 0.001), while prior outcomes increased decision latency (b = 31.50, *p* < 0.001). Following inspection of Fig 3C, a *post hoc* model revealed an interaction between cues and previous outcomes (b = 17.36, *p* < 0.05), indicating that in the presence of cues, previous outcomes further increased decision latency (Table 1).

### fMRI – Whole-brain Analysis

An exploratory whole-brain multivariate ANOVA (Table S1) was conducted to test for effects of cue condition (cued, uncued), choice type (risky vs safe), outcome (win vs non-win), and VGT phase (decision, anticipation, and feedback). Although outcome information was not available to participants at the decision and anticipation phases, this omnibus model was fully crossed on the assumption that these interactions would average out to zero. Significant main effects were observed only for phase, with large clusters showing differences across the three phases in primary auditory cortex (A1), large sections of visual cortex (V1), frontoparietal networks, dorsolateral and ventrolateral prefrontal cortices, but also subcortical networks including the basal ganglia, bilateral anterior insula (AntIns), and smaller clusters in OFC. *Post hoc* generalized linear t-tests indicated greater activity for feedback, compared to decision and anticipation phases, in a large OFC cluster, dorsomedial prefrontal cortex, as well as left auditory cortex, precuneus, bilateral lateral intraparietal/supramarginal gyrus. Decision and anticipation phases showed greater activity compared to feedback in frontoparietal regions including motor regions and lateral prefrontal cortex, visual association regions including V1, and networks related to motivation including posterior cingulate cortex, bilateral AntIns, basal ganglia, and the midbrain.

Further differences were observed between the anticipation and decision phases. Compared to anticipation, the decision phase elicited greater activity in the frontoparietal network, visual association regions, dorsolateral prefrontal, insula, posterior cingulate, a small OFC cluster, bilateral putamen, and midbrain areas. Anticipation only elicited greater activity than the decision phase in caudate, and bilateral superior temporal sulcus, near the auditory cortex. Some of these effects were driven in part by deactivation, for example, the OFC and nucleus accumbens (NAcc) effects in the decision > anticipation contrast were driven by lower activity than baseline during the anticipation phase, not increased activity during the decision phase (simple effects, Table S1).

The main effect of phase was qualified by phase x outcome and phase x cue condition interactions, with large sensory cortical clusters in bilateral auditory cortex and visual cortex seen in both cases. A large cluster in auditory cortex was also seen in the three-way interaction between outcome x cue condition x phase, given that the augmented auditory and visual feedback were only present in the cued condition, and when participants won. Additional noteworthy clusters were seen in the outcome x cue condition x phase interaction, including a small AntIns cluster and a cluster just ventral of the NAcc (see Fig 4). No cluster survived FDR correction in any other two or three-way interaction, or in the full four-way interaction.

**Figure 4.**
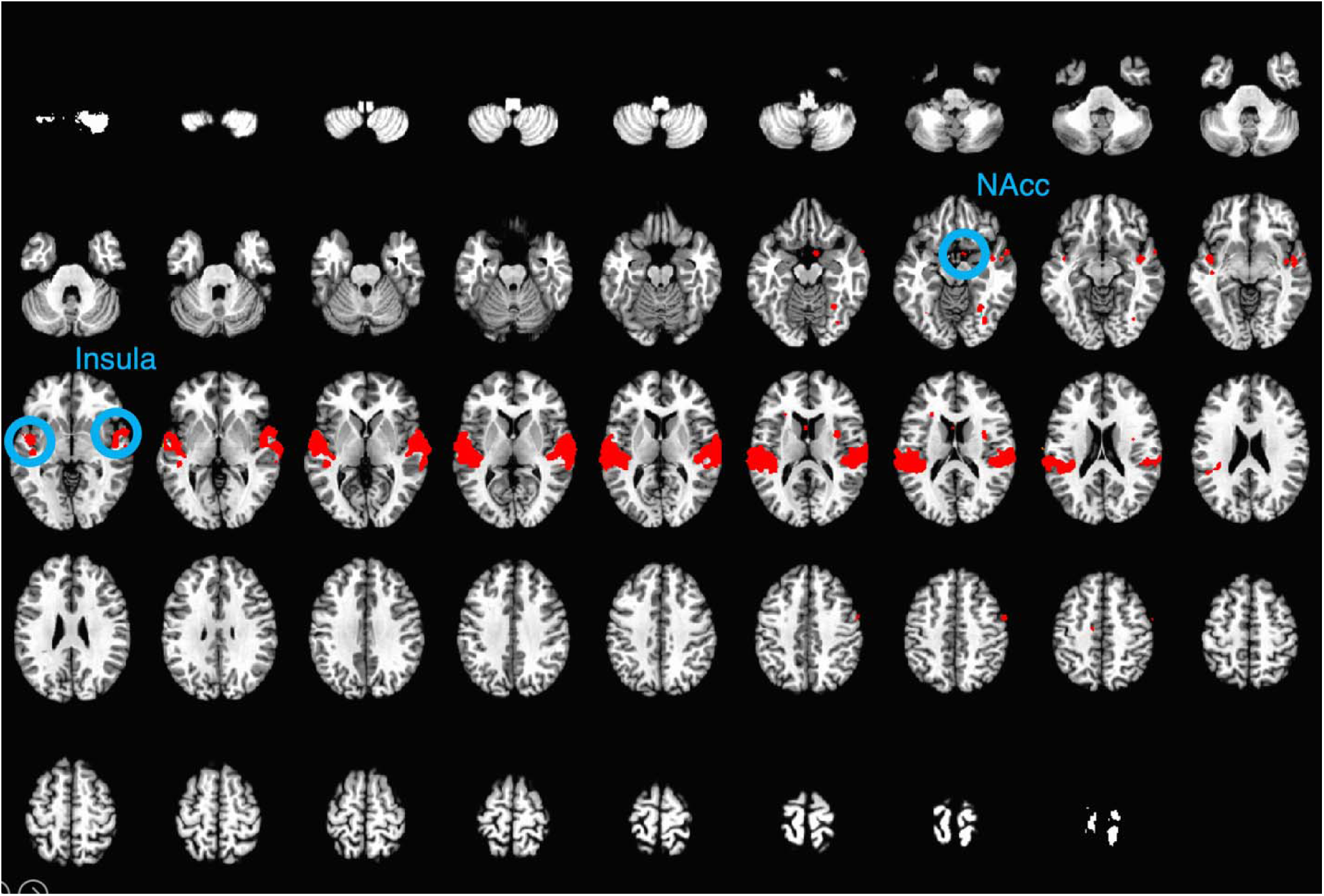
Whole-brain outcome x cue x phase effects. This F contrast shows prominent bilateral insula clusters and bilateral temporal clusters, and a small nucleus accumbens cluster (FDR *q* = 0.05, k = 5).

### Region of Interest (ROI) Analysis

To test whether the cue condition and risky choice modulated reward-related regions during decision and anticipation phases, linear mixed models were conducted on extracted signal from the NAcc and OFC ROIs. Participants were entered as random effects, and cue condition and risky choice as fixed effects. Binary regressors were effect coded. In the decision phase, choosing the risky option increased NAcc activity (Table S2, b = 0.011, *p* < 0.02) compared to the grand mean. During anticipation, NAcc activity was higher before a risky outcome compared to the grand mean (Table S2, b = 0.015, *p* < 0.002). In the decision phase, OFC activity was greater than the grand mean on risky choices (Table S3, b = 0.015, *p* < 0.05), but this effect was not observed during anticipation.

In line with the behavioral analyses, we conducted additional analyses to examine parametric effects with EVR, and the outcome of the previous trial. No effects were observed in the EVR model. Modelling win outcomes on the previous trial, risky choices increased NAcc signal at decision (b = 0.018, *p* = 0.01), as did winning outcomes on the previous trial (b = 0.010, *p* < 0.05). These effects were qualified by a three-way interaction between risk, cue condition, and last trial outcome (b = −0.010, *p* < 0.05), such that the increases in NAcc signal caused by choosing the risky option and from prior outcomes were dampened in the presence of cues.

We next determined whether cue condition, choice type, and outcome impacted ROI activity at the time of feedback. For NAcc, winning outcomes increased signal at feedback (b = 0.018, *p* < 0.001) in a basic model including only outcome and cue condition. When choice type was included (Table S4), the effect of outcome persisted (b = 0.020, *p* < 0.001), and was qualified by an interaction with risky choice (b = 0.010, *p* = 0.01): NAcc signal was augmented by risky wins (Fig 5A). We did not observe any significant main effects or interaction terms in the OFC ROI (Table S5). Our pre-registered hypotheses, that activity in the NAcc and OFC would be modulated by cue condition, were therefore not supported. In analyses of AntIns activity, a significant effect was observed in the three-way interaction between risk, win outcome, and cues, such that when participants won following risky choices in the cued condition, AntIns signal increased (b = 0.009, *p* < 0.01, Fig 5C).

**Figure 5.**
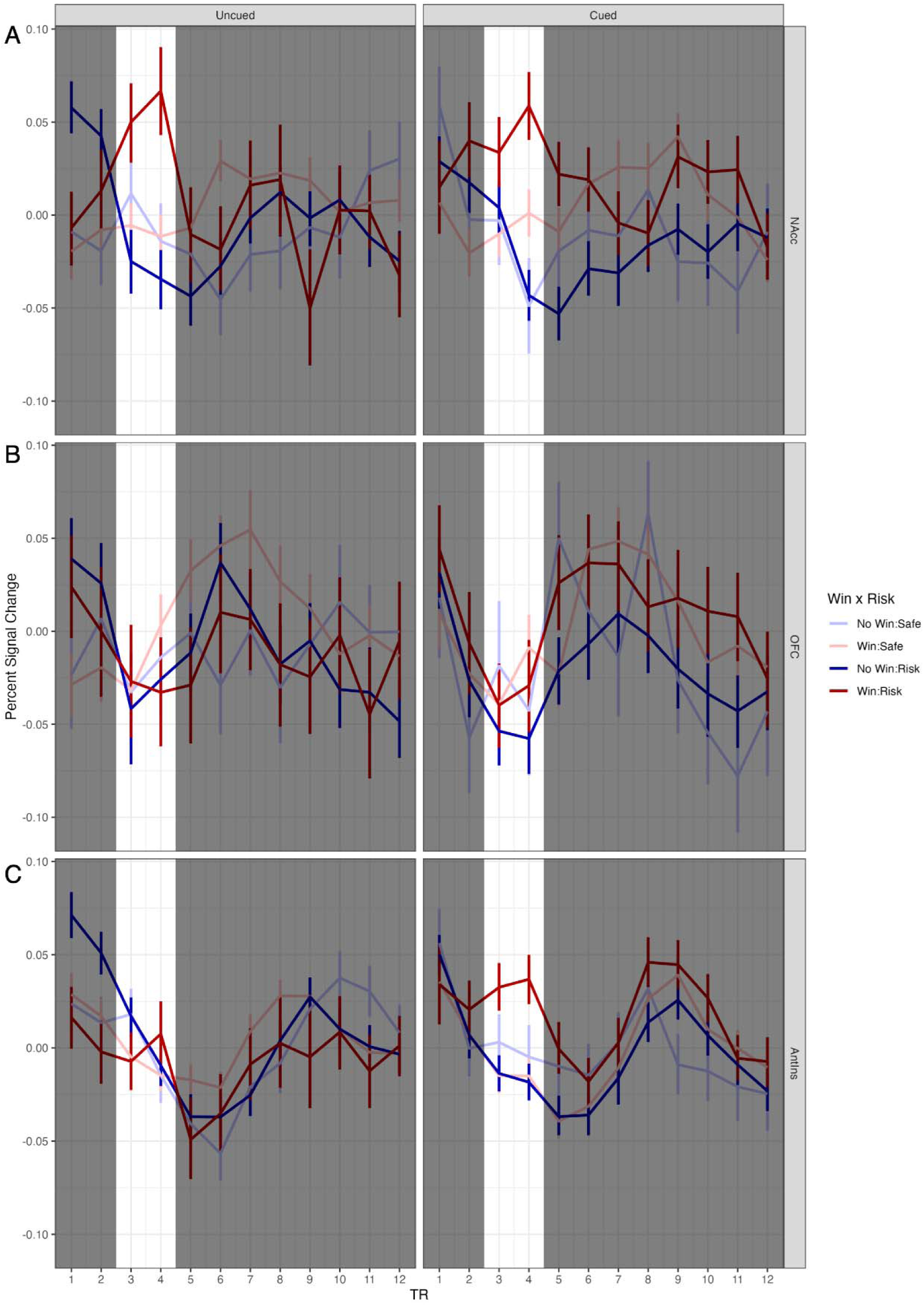
ROI signal change at feedback as a function of choice type and trial outcome. (A) Winning outcomes evoked greater NAcc activity than the grand average. Risky choice augmented this effect, as indicated by a significant choice type x outcome interaction. (B) Activity within the OFC did not change significantly based on outcome or cue condition (C) An exploratory analysis revealed a three-way interaction between cue condition, choice type, and trial outcome, indicating greater insula activity specifically for cued risky wins.

## Discussion

The present study replicates previous behavioral results showing that the incorporation of audiovisual cues into a two-choice lottery task increases risky choice (Cherkasova et al., 2018; Cherkasova et al., 2024), in an independent sample. The task modifications extend this observation to an inter-leaved block design appropriate for fMRI. Neural activity during the decision phase was greater within both the NAcc and OFC when participants made risky choices. NAcc activity was also higher during anticipation following risky choices, and in response to winning outcomes, and especially in response to risky wins. Unexpectedly, neither the NAcc nor OFC activity differentiated between cued and uncued trials across any task phase, thus failing to support our pre-registered hypotheses for our two *a priori* ROIs. Exploratory analyses revealed that activity within the AntIns increased in response to risky wins, in the cued condition only. Signaling within the AntIns may therefore contribute to the ability of win-paired audiovisual cues to potentiate risky decision making.

Decision latencies were also sensitive to the cue manipulation, being longer on cued trials and particularly following winning outcomes. In a recent study, adding audiovisual cues slowed choice latencies on a slot machine task, while simultaneously reducing the time participants spent looking at probability information (Baumann et al., 2025). Rats that made riskier choices when cues were present in the rGT were also slower to initiate the next trial after a cued win (Hales, Hrelja, et al., 2025). Slower response times following wins – the ‘post reinforcement pause’ (PRP) (Delfabbro & Winefield 1999) - are generally interpreted as reflecting the consummatory ‘savoring’ of positive outcomes, and the magnitude of this effect in a more realistic slot machine task correlated with game enjoyment (Dixon et al., 2019). People also prefer playing slot machines that incorporate winning sounds (Dixon et al., 2010). In this context, we note the latencies on the VGT are considerably slower than the operant-level responses commonly used to detect PRPs, and it remains unclear whether this slowing reflects comparable mechanisms. We also cannot parse in the current design whether the differences as a function of the previous trial outcome are driven by post-loss speeding (i.e. frustration or loss-induced impulsivity) rather than post-win slowing (Dyson, 2024; Eben et al., 2020). Nevertheless, these data complement recent work showing that the addition of win-paired audiovisual cues increase decision latencies as well as risky choice *per se*.

If participants interpreted cued wins as more valuable than uncued wins, as might be inferred from the latency data, one might predict that activity should be relatively greater in brain regions sensitive to reward on cued trials. Reward size and risk cannot be separated in the current design, in that the larger reward on each gamble is always conferred by the more risky choice. Both reward magnitude and uncertainty can elevate NAcc activity (Bartra et al., 2013; Knutson et al., 2001; Kuhnen & Knutson, 2005; Matthews et al., 2004). In keeping with these reports, fMRI signal within the NAcc was greater during the decision phase before risky choices, when anticipating the outcome of a risky gamble, and when a risky reward was received at feedback, but these elevations were observed regardless of cue condition. Signal within the OFC ROI was also greater during the decision phase when participants made risky choices, but again did not differentiate between cue conditions. These null findings for our cue manipulation suggest that cue-induced risky choice may not be driven by the overvaluation of rewards, at least within these traditionally reward-associated brain regions. Thus, the neural locus through which win-paired cues promote risky decision-making may lie elsewhere.

The main evidence for differential activity between cued and uncued trials emerged in the AntIns, wherein activity rose selectively during the feedback phase after cued, risky wins. Greater activity within the AntIns has been linked to motor impulsivity (Ramautar et al., 2006; Uddin, 2015), but it seems unlikely that the rise in AntIns activity during cued risky wins lead to impulsive responding here. Motor disinhibition in response to win-paired cues would arguably result in faster choices on cued trials, but as already discussed, participants exhibited longer choice latencies following wins during cued blocks. Using RL models, AntIns activity has been associated with negative reward prediction errors, such that activity increases when outcomes are worse than expected (Garrison et al., 2013). The VGT does not lend itself to RL modeling, but this signal could plausibly arise when subjects fail to win on safe (i.e. low risk) trials. We did not find this pattern in any region analyzed. Perhaps critically, participants did not *lose* money on the non-win trials, but simply failed to gain. Although reward-sensitive brain areas are thought to process such zero point outcomes as losses (Nieuwenhuis et al., 2005), future studies using mixed gambles that include the risk of financial loss may provide stronger evidence for AntIns activity in relation to a reward prediction error framework.

The AntIns forms part of the frontoparietal salience network, and is probably best known for its role in interoceptive awareness (Craig, 2002, 2003, 2009; Critchley, 2005; Critchley et al., 2004; Seeley et al., 2007). The rise in AntIns activity upon receipt of cued risky wins may reflect physiological arousal to such events, which may impinge on our bodily awareness to a greater degree than smaller cued wins or uncued rewards. In a previous study, phasic changes in pupil diameter were larger during the decision and anticipation phases of the VGT when subjects were gambling for cued rewards, even though the graphics presented during these trial epochs were identical (Cherkasova et al., 2018). As such, win-paired cues may increase physiological arousal during game play. The rush and thrill associated with probabilistic wins is a recognized driver of gambling’s appeal (Baudinet & Blaszczynski, 2013), and it is plausible that the AntIns signal at time of feedback contributes to the sense of excitement resulting from a cued risky win.

AntIns activity has also been described when participants made “slips of action” following a contingency change on an operant learning task, i.e. when *habitual* errors were committed (van Timmeren et al., 2023). In rats, adding win-paired cues to the rGT renders decision-making insensitive to reinforcer devaluation, indicating choice is no longer truly goal-directed but at least partially controlled by the habit system (Hathaway et al., 2022; Hathaway et al., 2021). AntIns lesions reduced sign-tracking in rats (Pribut et al., 2022), a behaviour that is thought to reflect habitual control (Morrison et al., 2015). Eye-tracking data suggest human subjects spend relatively less time looking at the graphics containing information about outcome probability when the ensuing rewards are cued, perhaps leading to less-informed or less model-based choice (Baumann et al., 2025; Cherkasova et al., 2018). Together, these results may suggest that AntIns activation in response to large, cued, risky wins facilitates a reduction in goal-directed control over choice, leading to the habitual selection of more risky options. While this hypothesis is clearly speculative at present, it may warrant further investigation.

Insula damage can a result in the cessation of nicotine dependence (Naqvi et al., 2007), and considerable data have linked activity in this region to the overwhelming urge to use drug, also known as craving (Droutman et al., 2015; Murray et al., 2025). In animal studies, silencing the AntIns disrupts the ability of drug-paired cues and contexts to induce drug-seeking (Agoitia et al., 2024; Arguello et al., 2017). Gambling cues can themselves induce the urge to gamble, and the degree of craving experienced correlates positively with AntIns activity (Limbrick-Oldfield et al., 2017). The current finding that cued, risky wins increased AntIns activity in healthy volunteers may point to a mechanism by which cue-induced craving develops in those vulnerable to gambling disorder. Future work evaluating how this AntIns response changes with gambling experience, particularly in those experiencing harms from gambling, would be helpful in substantiating this hypothesis.

As more of our lives are lived on-line, we are increasingly exposed to games and apps which make heavy use of reward-concurrent audiovisual cues. Identifying brain regions that are differentially activated by the presence of such stimuli can offer insight into how these cues alter cognition and affect. To our knowledge, this is the first neuroimaging study to explicitly investigate how adding “bells and whistles” to a laboratory-based gambling task alters patterns of brain activation. Our observation that cued, risky wins selectively activate the AntIns may ultimately inform strategies to combat the risk-promoting effects of win-paired cues, and mitigate their deleterious effects in those struggling with gambling dependency.

## Declarations

## Acknowledgements

This work took place at a UBC campus situated on the traditional, ancestral, and unceded land of the x məθk əy əm (Musqueam) People. We acknowledge and are grateful for their stewardship of this land for thousands of years.

## Availability of data, materials, and analysis code

Data, materials, and analysis code are available from the authors upon request. Our hypotheses and analysis plan were pre-registered (https://doi.org/10.17605/OSF.IO/J6SE4)

## Consent to participate

Participants gave written informed consent.

## Ethics approval

The study was conducted in accordance with institutional guidelines and the Declaration of Helsinki, and was approved by the Research Ethics Board of the University of British Columbia (H22-01101).

