## Supplemental material for "Anterior insula activity increased by cued risky wins in healthy volunteers"

**Table S1.** Exploratory omnibus whole brain analysis. (k=5; q<0.05)

| **Contrast**  Region | Volume  (Voxels) | Peak F-statistic | | Peak coordinate (Talairach) | | | |
| --- | --- | --- | --- | --- | --- | --- | --- |
|  |  | |  | | *x* | *y* | *z* |
| **Phase** |  | |  | |  |  |  |
| Bilateral frontoparietal, temporal, occipital, and subcortical regions. Anterior cingulate cortex, orbitofrontal cortex, caudate, precentral gyrus, superior parietal lobule, intraparietal lobule, V1, A1, cerebellum | 34034 | | 88.57 | | 4.5 | -7.5 | 44.5 |
| L middle temporal gyrus | 22 | | 8.14 | | 67.5 | 16.5 | -12.5 |
| R inferior temporal gyrus | 17 | | 12.41 | | -52.5 | 1.5 | -36.5 |
| R superior frontal gyrus | 14 | | 8.33 | | -16.5 | -61.5 | 23.5 |
| L inferior frontal gyrus | 11 | | 6.82 | | 49.5 | -25.5 | 11.5 |
| R inferior frontal gyrus | 7 | | 8.45 | | -58.5 | -25.5 | 17.5 |
| R inferior temporal gyrus | 6 | | 10.48 | | -49.5 | 10.5 | -24.5 |
| Orbitofrontal cortex | 6 | | 5.88 | | -1.5 | -22.5 | -21.5 |
| L superior frontal gyrus | 6 | | 6.10 | | 10.5 | -58.5 | 20.5 |
| **Outcome x Phase** |  | |  | |  |  |  |
| R superior temporal gyrus (A1) | 50 | | 44.75 | | -64.5 | 19.5 | 11.5 |
| L superior temporal gyrus (A1) | 30 | | 43.75 | | 46.5 | 10.5 | 5.5 |
| R superior temporal sulcus | 22 | | 44.24 | | -58.5 | 10.5 | 2.5 |
| R superior temporal gyrus | 13 | | 32.49 | | -52.5 | -1.5 | -3.5 |
| R posterior superior temporal gyrus | 8 | | 37.66 | | -52.5 | 31.5 | 14.5 |
| L superior temporal gyrus | 5 | | 20.95 | | 49.5 | 28.5 | 11.5 |
| **Cues x Phase** |  | |  | |  |  |  |
| R superior temporal gyrus (A1) | 396 | | 44.59 | | -61.5 | 10.5 | 2.5 |
| L superior temporal gyrus (A1) | 313 | | 58.21 | | 49.5 | 22.5 | 8.5 |
| L calcarine sulcus (V1) | 164 | | 24.03 | | 4.5 | 94.5 | 2.5 |
| R calcarine sulcus (V1) | 29 | | 19.31 | | -19.5 | 91.5 | 2.5 |
| R middle occipital gyrus | 6 | | 14.44 | | -31.5 | 88.5 | 8.5 |
| L lateral occipital | 5 | | 15.64 | | 22.5 | 91.5 | 17.5 |
| **Risk x Outcome x Phase** |  | |  | |  |  |  |
| Supplementary motor area | 30 | | 23.91 | | -4.5 | -10.5 | 53.5 |
| Posterior cingulate cortex | 10 | | 18.43 | | 4.5 | 43.5 | 23.5 |
| R thalamus | 5 | | 18.12 | | -10.5 | 25.5 | 2.5 |
| Thalamus | 5 | | 18.42 | | 1.5 | 22.5 | 5.5 |
| Superior medial frontal gyrus | 5 | | 18.10 | | -4.5 | -37.5 | 38.5 |
| **Outcome x Cues x Phase** |  | |  | |  |  |  |
| L superior temporal gyrus (A1) | 525 | | 62.19 | | 49.5 | 16.5 | 11.5 |
| R superior temporal gyrus (A1) | 453 | | 52.99 | | -55.5 | 13.5 | 5.5 |
| R fusiform gyrus | 13 | | 23.67 | | -28.5 | 55.5 | -12.5 |
| R insula | 13 | | 19.63 | | -31.5 | -1.5 | 14.5 |
| R frontal eye field | 9 | | 13.38 | | -55.5 | 4.5 | 44.5 |
| R ventral accumbens area | 8 | | 16.22 | | -7.5 | -1.5 | -12.5 |
| R fusiform gyrus | 7 | | 15.25 | | -31.5 | 73.5 | -12.5 |
| L fusiform gyrus | 5 | | 10.78 | | 31.5 | 61.5 | -12.5 |
| L caudate | 5 | | 14.74 | | 1.5 | -7.5 | 11.5 |
| L anterior insula | 5 | | 18.31 | | 25.5 | -22.5 | 14.5 |
| L middle cingulate cortex | 5 | | 15.53 | | 10.5 | 13.5 | 47.5 |

**Table S2. NAcc decision and anticipation regression results.** Statistics are coefficients and SEs. Significance (two-tailed) is indicated by **p* < 0.05; ***p*  < 0.01; ****p* < 0.001; ^†^*p* < 0.10 (nonsignificant trend).

|  | NAcc Activity (Decision) | | | NAcc Activity (Anticipation) | | |
| --- | --- | --- | --- | --- | --- | --- |
|  | Choice | Cues | Combined | Choice | Cues | Combined |
| Constant | 0.012  (0.006)* | 0.011  (0.006)^†^ | 0.012  (0.006)* | 0.015  (0.07)* | 0.013  (0.07)* | 0.015  (0.07)* |
| Risk | 0.011  (0.005)* |  | 0.011  (0.005)* | 0.015  (0.005)** |  | 0.014  (0.005)** |
| Cues |  | 0.001  (0.005) | -0.000  (0.005) |  | 0.006  (0.004) | 0.004  (0.004) |
| Risk x Cues |  |  | -0.004  (0.005) |  |  | -0.005  (0.004) |
| R^2^ | 0.0015 | 0.0000 | 0.0017 | 0.0026 | 0.0004 | 0.0032 |
| AIC | 1284.09 | 1289.67 | 1305.18 | 1453.35 | 1462.25 | 1472.78 |

**Table S3. OFC Decision and Anticipation Regression Results.** Statistics are coefficients and SEs. Significance (two-tailed) is indicated by **p* < 0.05; ***p* < 0.01; ****p* < 0.001; ^†^*p* < 0.10 (nonsignificant trend).

|  | OFC Activity (Decision) | | | OFC Activity (Anticipation) | | |
| --- | --- | --- | --- | --- | --- | --- |
|  | Choice | Cues | Combined | Choice | Cues | Combined |
| Constant | -0.001  (0.009) | -0.001  (0.009) | 0.001  (0.009) | 0.013  (0.011) | 0.011  (0.011) | 0.013  (0.011) |
| Risk | 0.010  (0.007) |  | 0.010  (0.007) | 0.015  (0.007)* |  | 0.015  (0.007)* |
| Cues |  | 0.000  (0.006) | -0.001  (0.007) |  | 0.010  (0.007) | 0.009  (0.007) |
| Risk x Cues |  |  | -0.008  (0.007) |  |  | -0.002  (0.007) |
| R^2^ | 0.0006 | 0.0000 | 0.0010 | 0.0013 | 0.0006 | 0.0018 |
| AIC | 4643.41 | 4645.77 | 4662.21 | 4378.29 | 4380.90 | 4396.57 |

**Table S4. NAcc Feedback Regression Results.** Statistics are coefficients and SEs. Significance (two-tailed) is indicated by **p* < 0.05; ***p* < 0.01; ****p* < 0.001; ^†^*p* < 0.10 (nonsignificant trend).

|  | NAcc Activity (Feedback) | | | | | |
| --- | --- | --- | --- | --- | --- | --- |
|  | | Choice | Cues | Outcome | Choice x Cues | Combined |
| Constant | | -0.007  (0.005) | -0.007  (0.005) | -0.009  (0.005)^†^ | -0.007  (0.005) | -0.005  (0.005) |
| Risk | | -0.003  (0.004) |  |  | -0.003  (0.004) | 0.004  (0.004) |
| Cues | |  | 0.001  (0.004) |  | 0.001  (0.004) | 0.001  (0.004) |
| Outcome | |  |  | 0.018  (0.004)*** |  | 0.020  (0.004)*** |
| Risk x Cues | |  |  |  | 0.002  (0.004) | 0.002  (0.004) |
| Risk x Outcome | |  |  |  |  | 0.010  (0.004)* |
| Cues x Outcome | |  |  |  |  | 0.001  (0.004) |
| Risk x Cues x Outcome | |  |  |  |  | 0.001  (0.004) |
| R^2^ | | 0.0002 | 0.0000 | 0.0060 | 0.0003 | 0.0080 |
| AIC | | -294.121 | -293.42 | -318.24 | -271.78 | -259.09 |

**Table S5. OFC Feedback Regression Results.** Statistics are coefficients and SEs. Significance (two-tailed) is indicated by **p* < 0.05; ***p* < 0.01; ****p* < 0.001; ^†^*p* < 0.10 (nonsignificant trend).

|  | OFC Activity (Feedback) | | | | | |
| --- | --- | --- | --- | --- | --- | --- |
|  | | Choice | Cues | Outcome | Choice x Cues | Combined |
| Constant | | -0.009  (0.008) | -0.008  (0.008) | -0.009  (0.008) | -0.009  (0.008) | -0.009  (0.008) |
| Risk | | -0.010  (0.006)^†^ |  |  | -0.009  (0.006) | -0.007  (0.006) |
| Cues | |  | -0.005  (0.006) |  | -0.005  (0.006) | -0.000  (0.006) |
| Outcome | |  |  | 0.010  (0.006)^†^ |  | 0.006  (0.006) |
| Risk x Cues | |  |  |  | -0.000  (0.006) | -0.001  (0.006) |
| Risk x Outcome | |  |  |  |  | 0.001  (0.006) |
| Cues x Outcome | |  |  |  |  | 0.001  (0.006) |
| Risk x Cues x Outcome | |  |  |  |  | 0.011  (0.006)^†^ |
| R^2^ | | 0.0007 | 0.0002 | 0.0007 | 0.0009 | 0.0020 |
| AIC | | 3301.90 | 3303.93 | 3301.75 | 3322.27 | 3359.11 |

**Table S6 AntIns Decision and Anticipation Regression Results.** Statistics are coefficients and SEs. Significance (two-tailed) is indicated by **p* < 0.05; ***p* < 0.01; ****p* < 0.001; ^†^*p* < 0.10 (nonsignificant trend).

|  | AntIns Activity (Decision) | | | AntIns Activity (Anticipation) | | |
| --- | --- | --- | --- | --- | --- | --- |
|  | Choice | Cues | Combined | Choice | Cues | Combined |
| Constant | 0.027  (0.005)*** | 0.026  (0.005)*** | 0.027  (0.005)*** | 0.031  (0.005)*** | 0.031  (0.005)*** | 0.032  (0.005)*** |
| Risk | 0.006  (0.004) |  | 0.006  (0.004)^†^ | 0.003  (0.004) |  | 0.003  (0.004) |
| Cues |  | -0.003  (0.004) | -0.004  (0.004) |  | 0.003  (0.004) | 0.002  (0.004) |
| Risk x Cues |  |  | -0.004  (0.004) |  |  | -0.004  (0.004) |
| R^2^ | 0.0001 | 0.0001 | 0.0006 | 0.0007 | 0.0002 | 0.0012 |
| AIC | -279.34 | -277.30 | -258.64 | -391.17 | -391.14 | -370.14 |

**Table S7. AntIns Feedback Regression Results.** Statistics are coefficients and SEs. Significance (two-tailed) is indicated by **p* < 0.05; ***p* < 0.01; ****p* < 0.001; ^†^*p* < 0.10 (nonsignificant trend).

|  | AntIns Activity (Feedback) | | | | | |
| --- | --- | --- | --- | --- | --- | --- |
|  | | Choice | Cues | Outcome | Choice x Cues | Combined |
| Constant | | -0.017  (0.004)*** | -0.017  (0.004)*** | -0.017  (0.004)*** | -0.017  (0.004)*** | -0.015  (0.004)*** |
| Risk | | 0.002  (0.003) |  |  | 0.002  (0.003) | 0.003  (0.003) |
| Cues | |  | 0.001  (0.003) |  | 0.001  (0.003) | 0.004  (0.003) |
| Outcome | |  |  | 0.003  (0.003) |  | 0.003  (0.003) |
| Risk x Cues | |  |  |  | 0.002  (0.003) | 0.002  (0.003) |
| Risk x Outcome | |  |  |  |  | 0.006  (0.003)^†^ |
| Cues x Outcome | |  |  |  |  | 0.002  (0.003) |
| Risk x Cues x Outcome | |  |  |  |  | 0.009  (0.003)** |
| R^2^ | | 0.0001 | 0.0000 | 0.0002 | 0.0002 | 0.0037 |
| AIC | | -2096.63 | -2096.20 | -2097.19 | -2073.53 | -2041.18 |

**Table S8. Risky Shift Group AntIns Regression Results.** Statistics are coefficients and SEs. Significance (two-tailed) is indicated by **p* < 0.05; ***p* < 0.01; ****p* < 0.001; ^†^*p* < 0.10 (nonsignificant trend).

|  | AntIns Activity (Feedback) | | |
| --- | --- | --- | --- |
|  | | High Risky Shift | Low Risky Shift |
| Constant | | -0.011 (0.007) | -0.018 (0.004)*** |
| Risk | | -0.005 (0.005) | 0.009 (0.004)* |
| Cues | | 0.005 (0.005) | 0.004 (0.004) |
| Outcome | | 0.003 (0.005) | 0.009 (0.004)* |
| Risk x Cues | | 0.004 (0.005) | -0.001 (0.004) |
| Risk x Outcome | | 0.005 (0.005) | 0.006 (0.004) |
| Cues x Outcome | | 0.004 (0.005) | 0.001 (0.004) |
| Risk x Cues x Outcome | | 0.010 (0.005)^†^ | 0.010 (0.004)* |
| R^2^ | | 0.0035 | 0.0071 |
| AIC | | -700.19 | -1297.52 |
